# Characterization of genomic DNA of lactic acid bacteria for activation of plasmacytoid dendritic cells

**DOI:** 10.1101/307074

**Authors:** Akira Horie, Yasuyuki Tomita, Konomi Oshio, Daisuke Fujiwara, Toshio Fujii

## Abstract

*Lactococcus lactis* strain Plasma (LC-Plasma) possesses strong activity of stimulating plasmacytoid dendritic cells (pDCs) via the TLR9-Myd88 pathway. To reveal the effective genome structure for pDCs stimulatory activity, we performed an *in vitro* screening, using randomly selected DNA fragments from the LC-Plasma genome. The results showed that CpG motifs are necessary factor for active DNA fragment, but the copy number of CpG motifs did not show strong correlation to the pDCs stimulatory activity of DNA fragment. We also found that the G+C contents of DNA fragments have significant negative effects on pDCs stimulatory activity. We also performed bioinformatics analysis of genome of lactic acid bacteria (LAB) and investigated the relation between CpG copy number in the genome and pDCs stimulatory activity. We found that strains of lactic acid bacteria (LAB) with high copy number of CpG motifs in the low-G+C region of the genome had higher probability of having high pDCs stimulatory activity. Three species, *L.lactis* subsp. *lactis, Leuconostoc mesenteroides*, and *Pediococcus pentosaceus* were the typical examples of high pDCs stimulatory activity LAB.

**Importance:** This study provides a new perspective on the structure of DNA fragments that are able to activate pDCs via the TLR9-Myd88 pathway. The information from this study should be useful for designing new DNA fragments, including phosphodiesterbond-DNA oligomers containing CpG motifs and DNA-containing vaccines. This work also presented an *in silico* screening method for identifying bacterial species that are able to activate pDCs. Therefore, this study should be useful for providing data for the development of vaccine adjuvants and therapeutics for infectious and allergic diseases.

## Introduction

Immunomodulatory effects of lactic acid bacteria (LAB) have attracted growing attention over decades. Numerous animal studies and clinical studies have demonstrated that LAB have antiallergic acitivity (1) and antivirus activity (2, 3). The cell products of probiotics that are responsible for immunomodulation are largely not known but might involve modifications of some of the known Microbe Associated Molecular Patterns (MAMPs) such lipoteichoic acids (LTA), exopolysaccharaides, RNA, and DNA. Interestingly, several studies suggested that strength of immunomodulatory activities depends on the species and strains on LAB (4–6).

Plasmacytoid dendritic cells (pDCs), a subset of dendritic cells (DCs) are immune cells that have a crucial function in the body’s defense against viral infections (7, 8). The pDCs originate in the bone marrow from myeloid and lymphoid precursors and require fms-like kinase 3 (Flt3L) for development. The pDCs sense DNA and RNA viruses through toll-like receptor 9 (TLR9) and TLR7, respectively, and subsequently produce interferon-alpha (IFN-α) (9), which induces the expression of genes coding for antiviral proteins such as MxA, viperin, and 2’-5’-oligoadenylate synthase. Several recent studies have revealed that pathogenic bacteria such as *Staphylococcus aureus* (10–12), *Neisseria meningitidis, Haemophilus influenza* (12), and *Streptococcus pyrogenes* (13) are able to enhance IFN-α production in mice and humans. However, well-known probiotic LAB strains such as *Lactobacillus* and *Bifidobacterium* have not yet been reported to activate pDCs.

We previously found that a specific strain of LAB, LC-Plasma (synonym of *Lactococcus lactis* subsp. *lactis* JCM 5805) was able to stimulate murine pDCs to produce IFN-α (5). Oral administration of LC-Plasma was found to result in significant immunomodulatory activity and enhanced antiviral activity in mice and humans (14–17). We also found that the LC-plasma stimulate pDCs via TLR9-Myd 88 pathway (5). This suggested that CpG motifs of genome DNA was the main MAMPs for pDCs stimulation.

It is well known unmethylated CpG motifs of bacterial genome that is the ligand of TLR9 (18, 19). Firstly, 5’-GACGTC-3’, 5’-AGCGCT-3’, and 5’-AACGTT-3’ was identified as *effi*cient immunostimulatory oligonucleotide ISS-ODN (20) and following studies proved that CpG containing hexamers, known as CpG motifs are able to stimulate B cells (18), and pDCs (21, 22). Various types of CpG-motifs have demonstrated as potent immunostimulatory DNA sequences (23). Studies of ODNs with phophoorotioate backbones for clinical application revealed the key structure of ISS-ODNs. For example, Hartmann et al. studied the effect of base change inside and outside of hexamers on activation of B cells and NK cells (24). Lenert et al. studied the extended sequence preferences both on ISS-ODN and immuno-inhibitory ODN (INH-ODN) on B cells (25). It was proposed that 5’-RRCGYY-3’ and 5’-GTCGTT-3’ are optimal consensus sequences for B cell activation in mice and primate, respectively (18, 24). The ISS-ODN containing CpG motif for pDCs activation was lately identified (22). The structural preference for ODN to activate pDCs was distinctly different from the ODN for B cells. 5’-RRCGRYCGYY-3’, 5’-RYCGYRTCGYR-3’, and 5’-RYCGRY-3’ were the most efficiently activate pDCs.

Later, the phophoorothioate bonded oligonucleotides containing the B cell stimulating motifs designated as Class B ODN and the phophoorothioate bonded oligonucleotides containing the pDCs stimulating motifs designated as Class A ODN. The importance of poly-G sequences at the 5’ end, the 3’ end have also been demonstrated. Fewer studies are carried out on phosphodiested bond backbone (21, 26, 27) and particular ODNs with high activity was also proposed.

In addition, several reports suggested that more specific CpG motifs or even non-CpG sequences of LAB are critical for proliferation of B cell activity, including BL07 motifs in *Bifidobacterium longum* BB536 (28), OL-LB7 motifs in *Lactobacillus delbrueckii* (29), ID35 motifs in *Lactobacillus rhamnosus* GG (30), and OL-LG10 motif from *Lactobacillus gasseri* JCM 1131 (31).

In this study, we constructed a library of genomic DNA fragments of LC-Plasma and investigated the pDCs stimulatory activity of each fragments to identify the essential character for pDCs activation. As we expected, the CpG motif was necessary for active DNA fragments. However, we found that the total copy number of CpG motifs in each DNA fragment was not strongly correlated with its pDCs stimulatory activity and that the G+C content of a genome DNA fragment has a significant effect on its potential for pDCs activation. We also performed an *in silico* analysis of the copy number of CpG motifs in the genome LAB and found that the low-G+C region of the genome has significant impact on the pDCs stimulation.

## Results

### CpG motifs are necessary for pDCs stimulatory activity of DNA fragments from LC-Plasma

In order to confirm that the necessity of CpG motifs for pDCs stimulatory activity, we performed *in vitro* experiment using PCR fragments. Four CpG-rich genomic loci (R1 R2, R3, and R4), and 2 CpG-free genomic loci (F1 and F2) were selected from the LC-Plasma genome. Three or four different fragments of each loci were selected and PCR primers were designed. The length and the copy number of CpG motifs in each fragments are shown in Table S1. In total, twelve CpG-rich DNA fragments and 7 non-CpG fragments were amplified and subjected to pDCs stimulating assay. The IFN-α production of pDCs stimulated with these amplified fragments was shown in Fig. 1. Eleven of 12 CpG-rich DNA fragments strongly induced IFN-α production, while none of the CpG-free fragments induced IFN-α production. These results strongly suggested CpG motif is necessary for pDCs stimulation.

**Fig 1.**
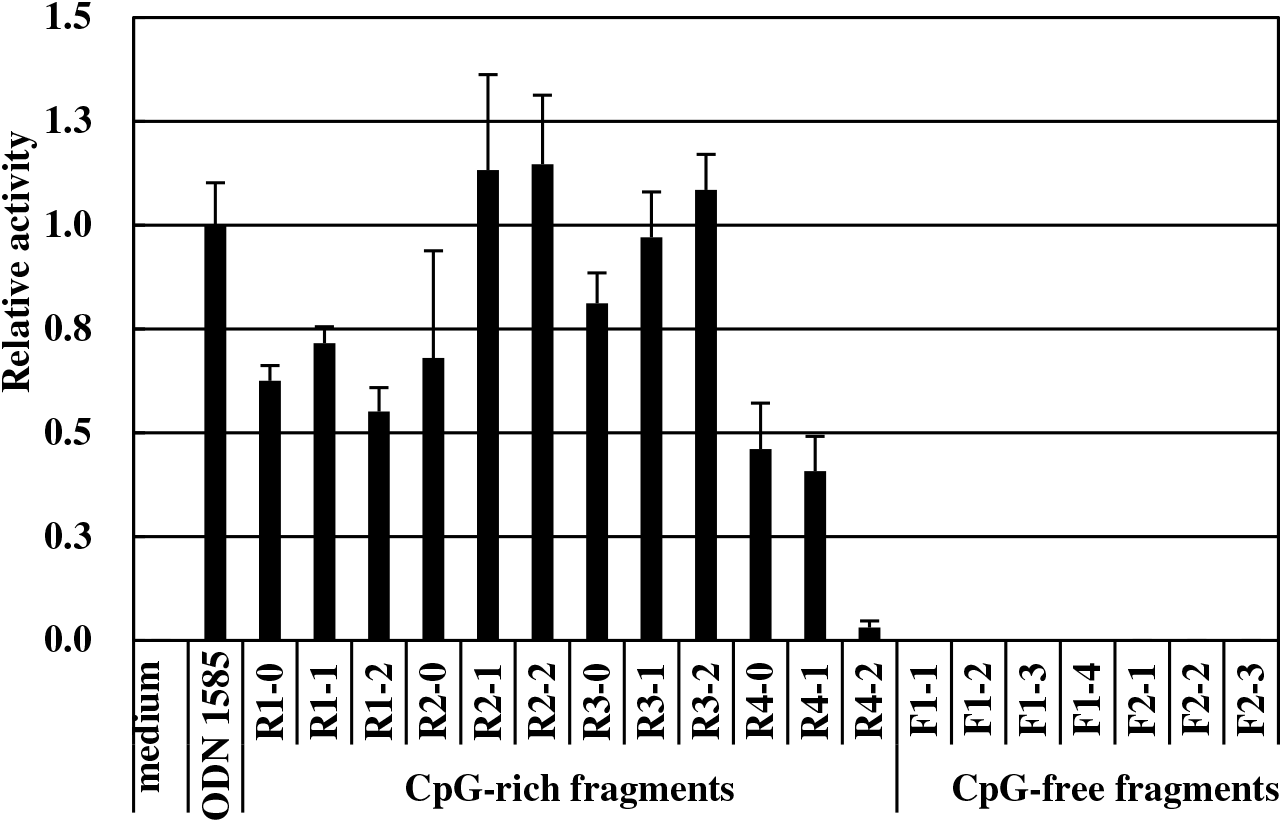
IFN-α induction by CpG-rich DNA fragments from *L.lactis* LC-Plasma. Flt3L-induced BM-DCs were stimulated by CpG-rich (R) or non-CpG (N) DNA fragments amplified from LC-Plasma genomic DNA. Each sample was added to cells at a final DNA concentration of 2 μg/mL Each value is the mean concentration ± S.D. for triplicate cultures.

### The copy number of CpG motifs are not strongly correlated to the pDCs stimulatory activity of DNA fragments from LC-Plasma

To reveal the copy numbers of CpG motif are related to the level of pDCs stimulatory activity, we constructed another library of DNA fragments from LC-Plasma. Fragments of approximately 200 bp with varied numbers of CpG motifs were randomly selected from the LC-Plasma genome (Table S2). The PCR-amplified fragments were subjected to assays for pDCs stimulatory activity.

We analyzed the correlation between pDCs stimulatory activity and copy number of CpG motifs in each DNA fragment (Fig. 2A). The results showed that the copy number of CpG motifs in the fragments was positively significantly correlated with activity (*p* < 0.01), and the correlation coefficient was *R*= 0.491, “moderate coefficient” defined by Guilford et.al. However, determination coefficient (*R^2^*) was only 0.24 which means another factor affects the pDCs stimulatory activity.

**Fig 2.**
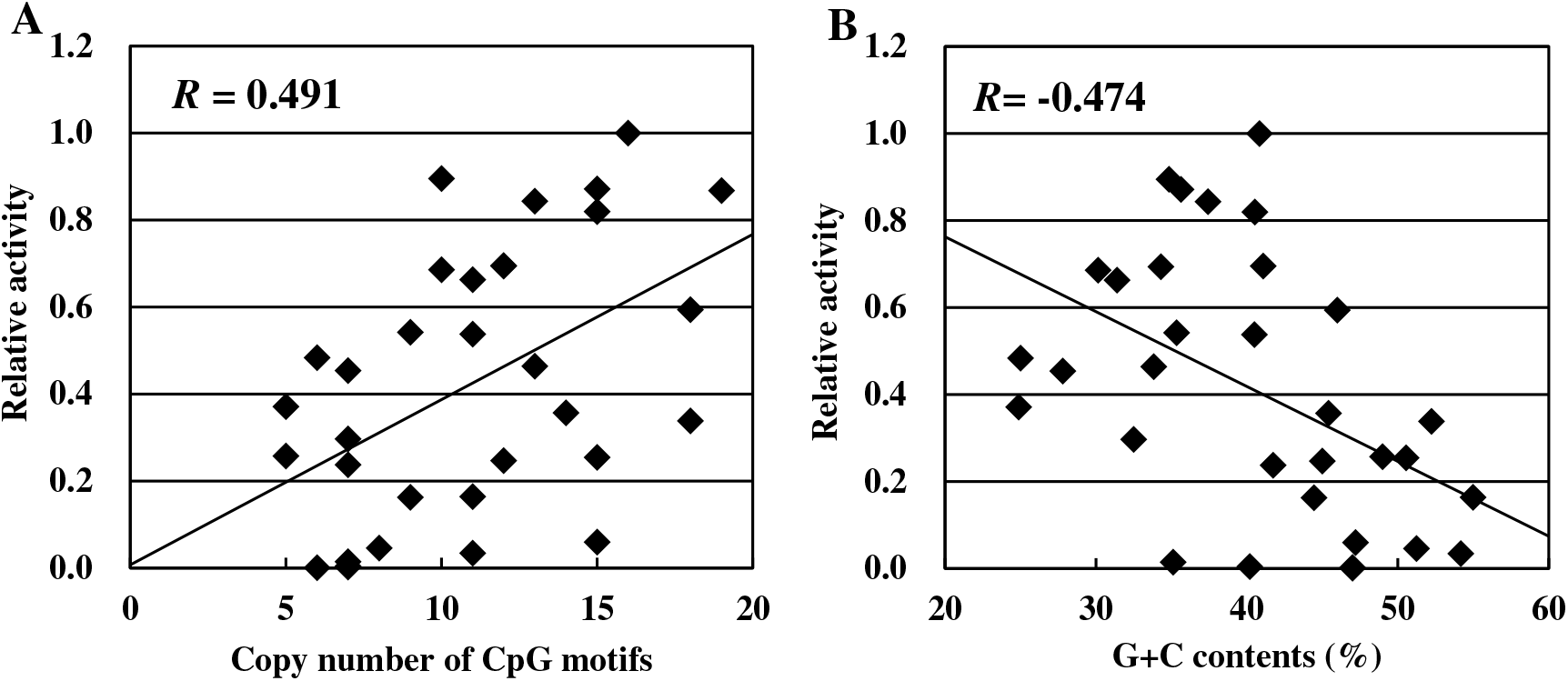
Correlation between immunostimulatory activity and the numbers of CpG motifs or G+C content contained in DNA fragments. Each dot depicts an independent 200 bp DNA fragment amplified from the LC-Plasma genome. Horizontal axes indicate A) the number of CpG motifs or B) G+C content contained in each DNA fragment. Vertical axes indicate the amount of IFN-α produced by BM-DCs stimulated by each type of DNA fragment.

### G+C content of DNA fragments from LC-Plasma is negatively correlated with pDCs stimulatory activity

We then studied the relation of the G+C contents of DNA fragments with the level of pDCs stimulatory activity. A significant negative correlation between pDCs stimulatory activity and G+C contents of the fragment (*R* = -0.474, *p* < 0.01, Fig. 2B) was observed. We performed bilayer stratified analysis based on G+C contents and compared the relation between the copy number CpG motifs and pDCs stimulatory activity. The DNA fragments into the low-G+C group composed of fragments with G+C < 40%, and the high-G+C group composed of fragments with G+C ≥ 40%. (Fig. 3A and 3B). The correlation coefficient was increased in both of the low-G+C group (*R* = 0.680, *p* < 0.01) and the high-G+C group (*R* = 0.647, *p* < 0.01). The degree of pDCs stimulatory activity per copy of CpG motifs was higher in the low-G+C group.

**Fig 3.**
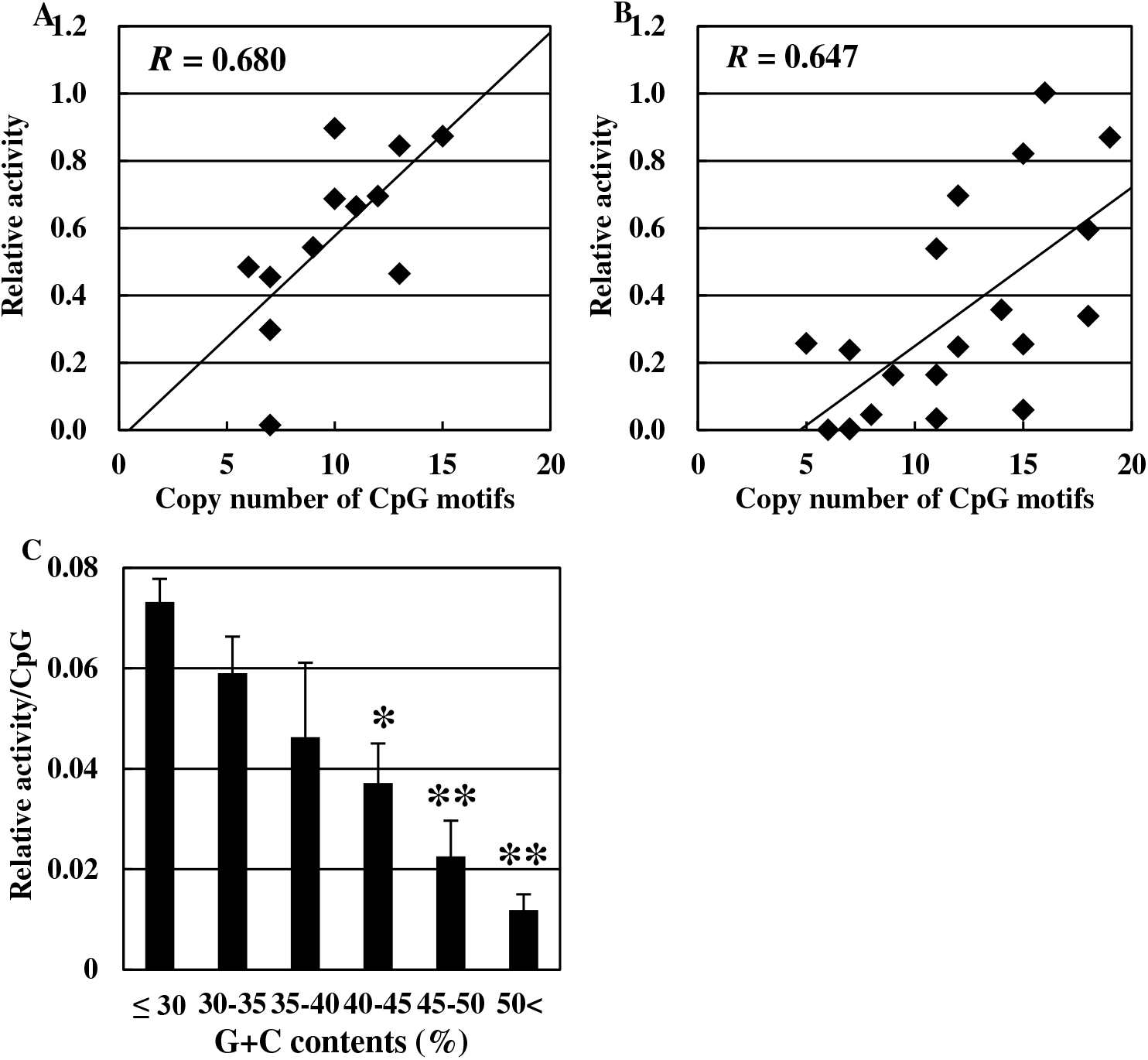
Correlation between the copy numbers of CpG motifs and IFN-α production in 200 bp DNA fragments. Each dot depicts an independent 200 bp amplified DNA fragment. A) amplified from the LC-Plasma genomic regions with G+C < 40%. B) amplified from the LC-Plasma genomic regions with G+C ≥ 40%. C) IFN-a production activities per CpG motifs in the DNA fragment having various G+C contents. IFN-a production by BM-DCs stimulated with each 200 bp DNA fragment was measured, and was divided by the copy numbers of CpG motifs. Bar depicts the standard deviation (S.D.). Bars with different notation exhibits significant difference (* *p*<0.05, ** *p*<0.01)

We also stratified DNA fragments into groups based on G+C contents as follows: < 30%, ≥30% to <35%, ≥35% to <40%, ≥40% to <45%, ≥45% to <50% and ≥50%. Stepwise reduction in pDCs stimulatory activity was observed, with a stepwise increase in G+C contents (Fig. 3C). We performed one-way ANOVA and Dunnet test. The results revealed that the levels of pDCs activity resulting from stimulation by fragments with G+C contents of ≥40% to <45%, ≥45% to <50%, and ≥50% were significantly lower compared to the activity induced by fragments with < 30% G+C. We also performed correlation analyses using randomly synthesized 300 bp fragments. Similar results were observed again (Fig. S1). These results strongly suggested that G+C content of DNA fragment is another essential factor to affect high level of pDCs stimulatory activity.

### Total copy number of CpG motifs in the genome DNA are not strongly correlated to the pDCs stimulatory activity of LC-Plasma

We carried out *in silico* analysis to investigate the relation between the copy number of CpG motifs and pDCs stimulatory activity. The total copy number of CpG motifs in the genome of *L. lactis* LC-Plasma was measured and compared to those of in the genomes of *Lactobacillus rhamnosus* ATCC 53103, and *Bifidobacterium longum* NCC 2705 which showed low pDCs stimulatory activity in a previous study (5). The results suggested that the number of CpG motifs in the LC-Plasma is three times smaller than that in the ATCC 53103 and four times smaller than that in NCC2705 (Table 1). We also measured the three of the pDCs-activating motifs, and two of B cell activating motifs in the genome of these LABs (Table 1). The results showed that the genome of ATCC 53103 contained 3.7 to 5.7 fold greater copy number of pDCs activating motifs and 1.7 to 5.7 fold greater copy number of B cells activating motifs than that of the genome of LC-Plasma. The genome of NCC2705 contained 5.6 to 17.4 fold greater copy number of pDCs activating motifs and 1.5 to 2.8 fold greater copy number of B cells activating motifs than that of the genome of LC-Plasma. These results suggested the copy number of CpG motifs are not strongly related to the level of pDCs stimulatory activity of LC-Plasma.

**Table 1.**
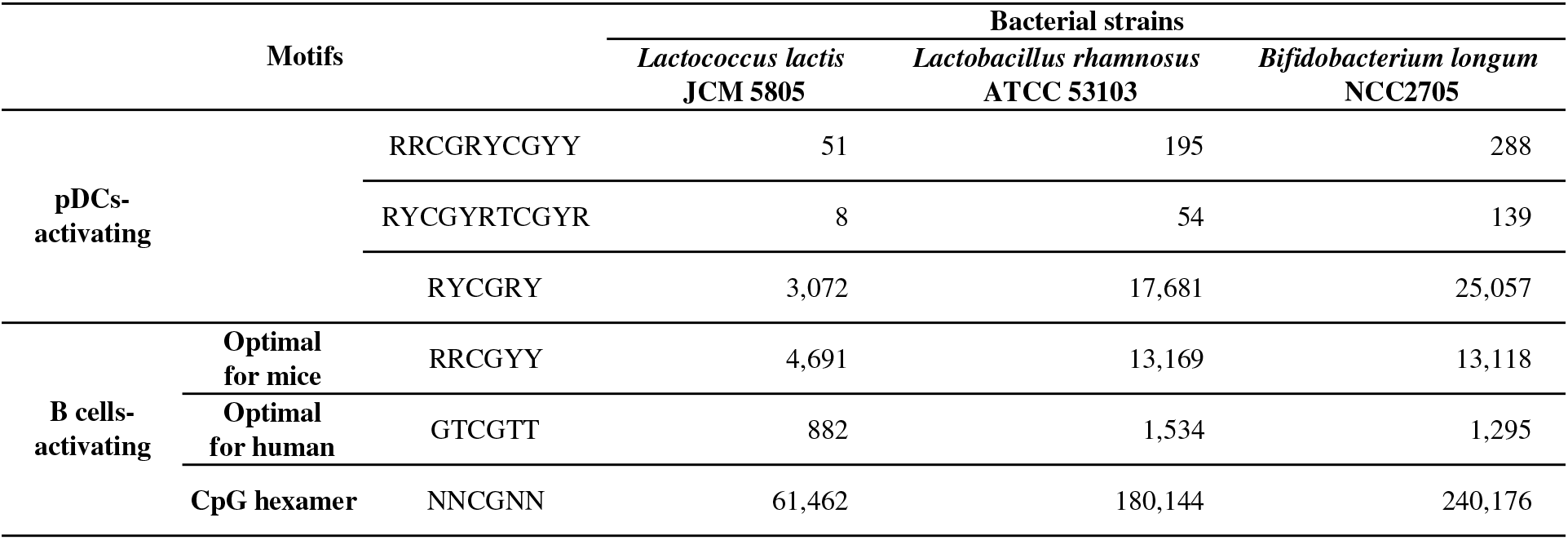
CpG motifs and copy numbers in the whole genome of LAB strains. Copy number of typical CpG motifs in the whole genome ofL. *lactis* LC-Plasma, *L. rhamnosus* ATCC 53103, and *Bifidobacterium longum* NCC2706 were determined as described in Materials and Methods.

### Comparing the pDCs stimulatory activity of single-stranded DNA

Because G+C content is directly related to the dissociation temperature of ds-DNA fragments, we evaluated pDCs stimulatory activity induced by synthetic oligonucleotides in single-stranded (ss) or double-stranded (ds) form. Two ss-CpG oligomers were synthesized, based on the sequences of ODN 1585 and ODN 2216 (InvivoGen, San Diego, CA, USA). As shown in Fig. 4, both oligonucleotides induced pDCs stimulatory activity, while their complementary sequences did not. We also synthesized the ds-form of ODN 1585 and ODN 2216, by annealing the normal and complementary strands. Interestingly, neither ODN 1585 nor ODN 2216 induced pDCs stimulatory activity in ds forms. In addition, the sense ODN hybridized with the antisense 6-bp sequence of the core CpG motif induced high pDCs stimulatory activity. These results suggest that an ss-CpG oligomer is more efficient at stimulating pDCs than a ds-CpG oligomer. The results also suggest that strong hybridization affinity between complementary strands might reduce the pDCs stimulatory activity of CpG motifs.

**Fig 4.**
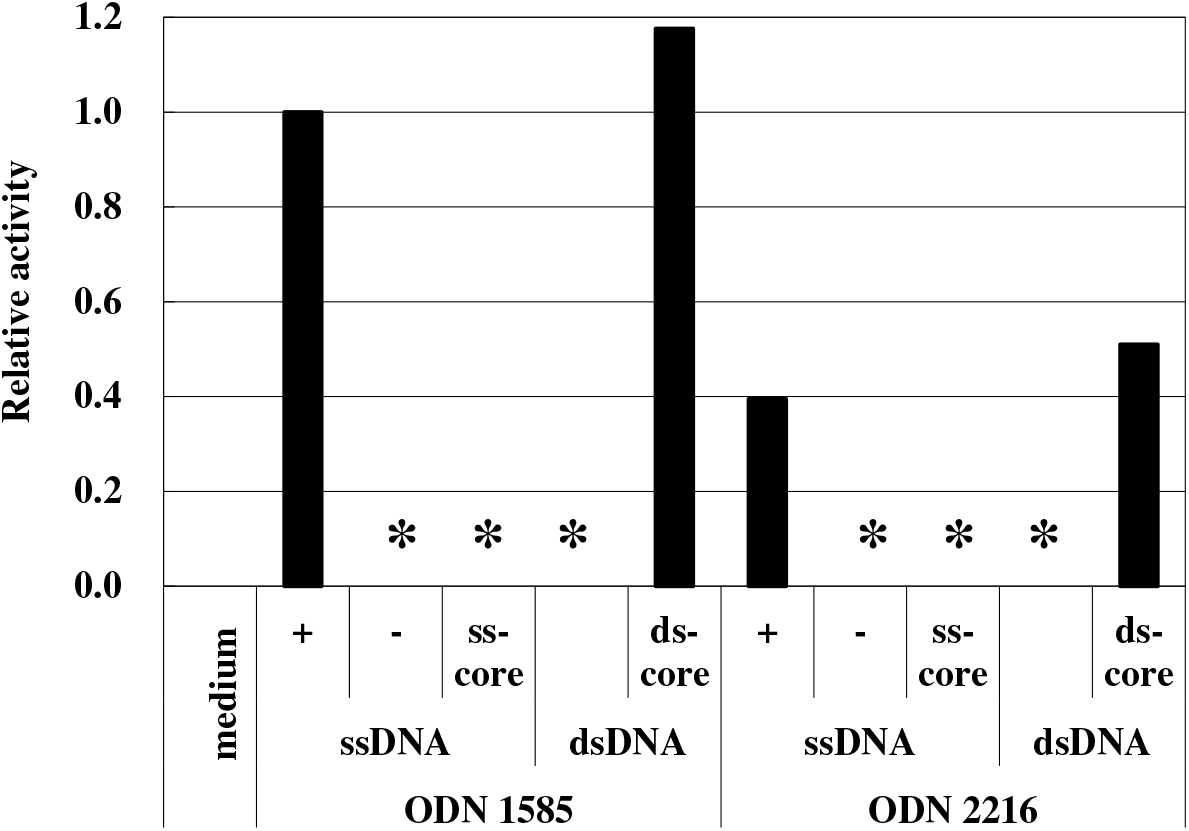
IFN-α production induced by single-or double-stranded forms of synthetic oligonucleotides. Single-stranded DNA oligomers (ssDNA) with phosphodiester bonds were synthesized, based on sequences of ODN1585 and ODN2216 (Invivogen, San Diego, CA, USA). Double-stranded DNA oligomers (dsDNA) were prepared by annealing sense and antisense strands of ssDNA. Each synthesized oligomer (2 μg) was tested on Flt3L induced BM-DCs, and the production of IFN-α was measured. + : sense strand; – : antisense strand; ss-core: 6-base CpG motif of sense strand; ds-core : Hybrid of sense strand DNA oligomer with antisense strand of 6 base CpG motif, *not detected

### *In silico* analysis of the copy number of CpG motifs in whole genome and low-G+C region of the genome of LAB

We investigated the frequency of CpG motifs in whole genomes and in the low-G+C region (<40% of G+C contents) of the genome (Fig. 5A). A linear increase of frequency of CpG motifs was observed with increasing G+C content of whole genomes. On the contrary, the frequency of CpG motifs localized to low-G+C regions of the genome showed an inverse correlation with the G+C content of whole genomes (Fig. 5B). Three species (*Lactococcus lactis* subsp. *lactis, Pediococcus pentosaceus*, and *Leuconostoc mesenteroides*) with the genomes of low G+C contents (35.2% to 37.7%) contains 20 copies/kb CpG motifs in their low-G+C regions, while the other four species (*L.plantarum, L.casei, L.fermentum*, and *Bifidobacterium longum*) with the genomes of high G+C contents (46.6% to 60.1%) contains less than 10 copies/kb CpG motifs in their low-G+C regions.

**Fig 5.**
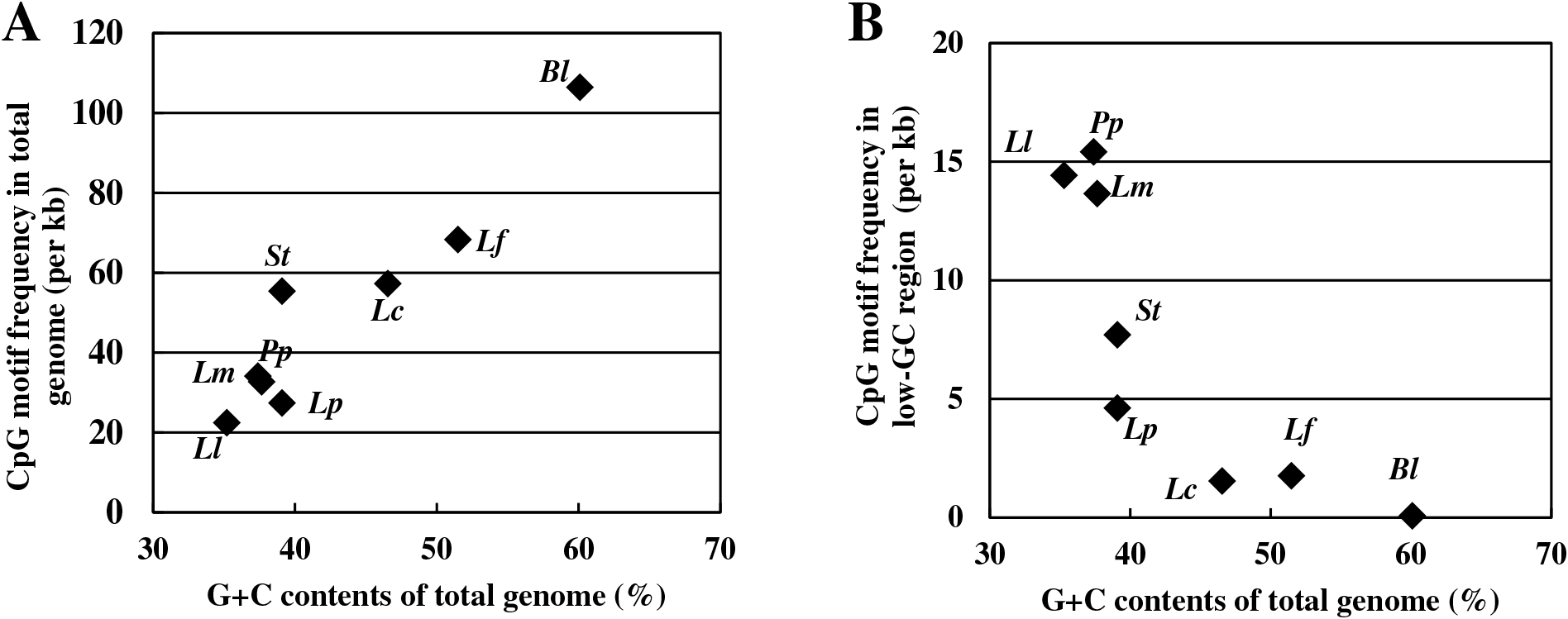
CpG motifs frequency in the genome of species of LAB. The frequency in each genome was depicted as a dot: A) whole genome and B) G+C < 40%. *Ll; Lactococcus lactis* LC-Plasma, *Pp*; *Pediococcus pentosaceus* ATCC 25745, *Lm; Leuconostoc mesenteroides* ATCC 8293, *St; Streptococcus thermophilus* CNRZ 1066, *Lp; Lactobacillus plantarum* WCFS1, *Lc; Lactobacillus casei* ATCC 334, *Lf; Lactobacillus fermentum* IFO 3956, *Bl; Bifidobacterium longum* NCC 2705

### The copy number of CpG motifs in the low-G+C region of the genome was closely related to the pDCs stimulatory activity of LAB

We investigated the differences of pDCs stimulatory activity between strains of these LAB species (Table 2). The wide variations of strains-based-activity were observed in each species. It was also observed that the frequencies of high-activity-strains were clearly different between the species. Five of 7 strains belonging to *L. lactis* subsp. *lactis* strains, two of 10 *L. mesenteroides* strains, and five of 19 *P. pentosaceus* strains induced marked (> 100 pg/mL) production of IFN-α. On the contrary, none of the LAB strains showing a lower frequency of CpG motifs in low-G+C regions, including *L. plantarum, L. casei*, and *L. fermentum*, exhibited significant stimulatory activity. The means of activity was also higher in the three strains compared to others. These results strongly suggest that the pDCs stimulatory activity of a bacterial strain depends on the copy number of CpG motifs in the low-G+C region of the genome and not on the copy number over the entire genome.

**Table 2.** pDCs stimulatory activity of LAB strains with varied GC contents in genome DNA. *Number of high (≥100 pg/mL)-IFN-α-producing strains per tested strains.

| Species | Strains | IFN-α<br>(pg/mL) | CpG<br>frequency (per kb)<sup>#</sup> | GC<br>(%) | High* |
| --- | --- | --- | --- | --- | --- |
| *Lactococcus lactis* | JCM 5805 | 404.4 |  |  |  |
|  | JCM 20101 | 389.8 | 22.3 | 35.3 | 2/3 |
|  | ATCC 15577 | 12.2 |  |  |  |
| | mean ± S.E. | 271.0 ± 24.6 | | | |
| *Pediococcus pentosaceus* | JCM 2026 | 84.0 |  |  |  |
|  | JCM 20109 | 353.3 |  |  |  |
|  | JCM 20314 | 102.1 |  |  |  |
|  | JCM 20459 | 5.5 |  |  |  |
|  | NBRC 3182 | 59.1 |  |  |  |
|  | NBRC 3891 | 15.8 |  |  |  |
|  | NBRC 3892 | 116.4 |  |  |  |
|  | NBRC 3893 | 32.2 |  |  |  |
|  | NBRC 3894 | 46.0 |  |  |  |
|  | NBRC 12229 | 18.7 | 28.4 | 37.4 | 5/19 |
|  | NBRC 12230 | 7.9 |  |  |  |
|  | NBRC 12232 | 14.8 |  |  |  |
|  | NBRC 12318 | 59.8 |  |  |  |
|  | NBRC 101982 | 404.4 |  |  |  |
|  | NBRC 101983 | 5.6 |  |  |  |
|  | NBRC 101984 | 5.9 |  |  |  |
|  | NBRC 101985 | 5.8 |  |  |  |
|  | NBRC 101986 | 172.3 |  |  |  |
|  | NBRC 101987 | 34.3 |  |  |  |
| | mean ± S.E. | 81.3 ± 114.5 | | | |
| *Leuconostoc mesenteroides* | NBRC 3349 | 3.2 |  |  |  |
|  | NBRC 3426 | 57.3 |  |  |  |
|  | NBRC 3832 | 353.5 |  |  |  |
|  | NBRC 12060 | 4.2 |  |  |  |
|  | NBRC 100495 | 5.7 |  |  |  |
|  | NBRC 100496 | 8.1 | 27.1 | 37.7 | 2/10 |
|  | NBRC 102497 | 36.0 |  |  |  |
|  | NBRC 102480 | 74.2 |  |  |  |
|  | NBRC 102481 | 167.0 |  |  |  |
|  | NBRC 107766 | 4.4 |  |  |  |
| | mean ± S.E. | 71.4 ± 111.6 | | | |
| *Lactobacillus acidophilus* | JCM 1132 | 11.7 | 15.4 | 34.7 |  |
|  | JCM 1551 | 16.7 |  |  |  |
| *Lactobacillus plantarum* | JCM 6651 | 55.2 | 4.6 | 39.1 | 0/5 |
|  | JCM 8341 | 4.0 |  |  |  |
|  | JCM 20110 | 9.8 |  |  |  |
| *Lactobacillus* low G+C | mean ± S.E. | 19.5 ± 20.5 | | < 40 | |
| *Lactobacillus casei* | ATCC 393 | 5.6 | 1.5 | 46.6 |  |
| *Lactobacillus rhamnosus* | ATCC 53103 | 3.9 |  | 47.0 |  |
| *Lactobacillus fermentum* | NBRC 3959 | 4.4 | 1.9 | 51.5 | 0/4 |
|  | NBRC 3961 | 5.2 |  |  |  |
| *Lactobacillus* high G+C | mean ± S.E. | 4.8 ± 0.8 | | 40 ≤ | |

We also carried out a statistical analysis of species-based pDCs stimulatory activity using Steel-Dwass method (Table S4). Significant differences were observed between *L.lactis* to *P. damnosus, L. mensteroides*, and *Lactobacillus* low G+C species. Marginally significant difference was also observed between *L. lactis* and *Lactobacillus* high G+C species. In addition, *P. pentosaceus* and *L. mesenteroides* also showed significant difference to *Lactobacillus* high G+C species.

## Discussion

Though stain differences of immunostimulatory activity of LAB has been the focus of interest for these decades, the molecular mechanisms has not yet been well understood. In the beginning of this study, we hypothesized that CpG copy number in the genome might be proportional to the pDCs stimulatory activity and that LC-Plasma may contain greater copy number of CpG motifs and/or some special sequences containing of CpG motifs. However, our results using DNA fragments and *in silico* analysis did not supported this hypothesis. In DNA fragment analysis, the CpG motifs seemed to be necessary for pDCs stimulation, but the correlation of the copy number of the CpG motifs and pDCs stimulatory activity was weak. In genome analysis, we could not find greater copy number of total CpG motifs, nor three of consensus sequences that have been reported as pDCs in the genome of LC-Plasma. In the long process of DNA fragment analysis, we found that the G+C contents have negative correlation of pDCs stimulatory activity. The stratification of DNA fragments based on G+C contents made the correlation between the copy number of CpG motifs and the pDCs stimulatory activity much stronger. In the genome analysis, we found that the CpG motifs in the low-G+C region of the genome is critical determinants of the pDCs stimulatory activity. Taken these together, we demonstrate that the G+C contents of DNA is one of the critical factor for pDCs stimulatory activity in either of DNA fragments or genome.

The effect of G+C contents on immunostimlatory activity has not yet been fully studied in the history of CpG motifs. Yamamoto et al. isolated DNA from bacteria, virus, invertebrate, vertebrate, and plant. They investigated the NK stimulatory activity of DNA samples but no correlation was observed between G+C contents and activity. To the best of our knowledge, this is the first study that has demonstrated that the G+C contents of DNA fragments has a direct effect on the immunomodulatory activity of pDCs.

Our results suggested that CpG fragment lost its pDCs stimulating activity by annealing to the complementary whole strand, while annealing of core sequence of CpG motif did not reduce the pDCs stimulating activity. This suggested that the dissociation is important for the CpG-motif containing DNA to stimulate pDCs. The CpG-motif containing DNA lost its activity by annealing the complementary strand. A recent study of crystal structures of 3 forms of TLR9 suggested that single-stranded oligonucleotides bound to TLR9 act as DNA agonists. It is possible that the G+C contents of DNA and affects the dissociation of ds-DNA fragments and the interaction with TLR9, which is followed by activation of pDCs. However, some investigators insist that oligonucleotides cannot occur in ss-forms (32), or that duplex structures are required for recognition by TLR9 (33, 34). It was suggested that the DNA sequence around the CpG motifs are also important for activation, since the fragment did not lost its activity by annealing the complementary strand of core CpG motif. Additional studies are needed to clarify whether single-strandedness is a key factor for pDCs activation.

Our results also presented a useful *in silico* screening methods of bacteria with high pDCs stimulatory activity at species level. We showed *L.lactis* subsp. *lactis, Pediococcus pentosaceus*, and *Leuconostoc mesenteroides* as typical example of higher pDCs stimulatory activity. In addition, our data also suggested that the copy number of CpG motifs in low-G+C region is not the absolute determinant of pDCs stimulatory activity of bacterial cells. We observed wide variety of pDCs stimulatory activities of strains in sole species. Moreover, the activity of *Lactococcus lactis* was significantly higher than those of *Pediococcus pentosaceus*, and *Leuconostoc mesenteroides*, though the copy number of CpG motifs in low-G+C region of those two species were higher than that of *Lactococcus lactis*. Not only the structure of DNA but also the bacterial cell’s affinity to pDCs or suitability of phagocytosis of pDCs affects the pDCs stimulating activity.

It should be noted that several previous studies were not consistent with our findings. Bioinformatics study by Kant *et. al*. suggested that the number of CpG motifs were highly negatively correlated with G+C content, which was also observed in this study. They also demonstrated that 5’-GTCGTT-3’ motif that is one of the most effective CpG motifs for humans had lower correlation with the G+C contents, and that the genomes of some probiotic strains had higher frequencies of 5’-GTCGTT-3’ motifs than intestinal bacterial strains (35). The observation in our study did not support their hypothesis since the frequency of 5’-GTCGTT-3’ motifs nor 5’-RRCGYY-3’ motifs did not correlate with the pDCs stimulating activity. Ménard *et al*. showed that CpG-rich DNA fragments with high G+C content from *Bifidobacterium longum* were effective for macrophage activation (36). We could not duplicate the findings of Ménard *et al* when we tested CpG-rich DNA fragments with high G+C content from *B. longum* on BM-derived DCs (data not shown). However, the effect of G+C contents may depend on host cell lineage. Singer *et al*. investigated the proportions of inflammation stimulatory (5’-RRCGYY-3’) and inhibitory (5’-NCCGNN-3’ and 5’-NNCGRN-3’) sequences in the genomes of pathogenic bacteria (37). They found species dependent differences in the proportion of stimulatory and inhibitory sequences, but they did not study the inflammatory responses of each pathogen. It would be a great interest whether the effect of G+C contents on inflammatory immune cytokines such as TNF-α, IL-1, and IL-6 caused by pathogenic bacteria.

In conclusion, our study provides a new perspective on the structure of DNA fragments that are able to activate pDCs via the TLR9-Myd88 pathway. Additional investigations and applications of our hypothesis may lead to the detailed understanding of host-bacterium interactions via TLR9 in other bacteria, other immune reactions, and other immunocytes.

## Materials and Methods

### Bacterial strains

The bacterial strains used in this study, *Lactococcus lactis* LC-Plasma and *Lactobacillus rhamnosus* ATCC 53103, were purchased from the collections held at the Japan Collection of Microorganisms (JCM) and American Type Culture Collection (ATCC), respectively. Other bacterial strains used in the screening assay were purchased from JCM, ATCC, or NITE Biological Resource Center (NBRC).

Cultures of bacterial strains were grown at 30°C or 37°C for 48 hours in De Man, Rogosa, and Sharpe (MRS) medium (BD Biosciences) or GAM medium (Nissui), which were prepared according to the suppliers’ instructions.

### Preparation of DNA fragments

Genomic DNA was extracted and purified from bacterial cultures grown as described in the previous section, using QIAGEN Genomic-tip 500/G (Qiagen). PCR amplifications of selected sequences, which were based on the results of our *in silico* analysis, were performed using the GeneAmp PCR System (Applied Biosystems), with primers designed according to the *L.lactis* LC-Plasma genome sequence. PCR was performed using TaKaRa Ex *Taq*^®^ (TaKaRa), according to the manufacturer’s instructions, using 10 ng of DNA template in 50 μl of reaction mixture containing primers at a concentration of 0.5 μM. The following thermal cycling profile was used: 5 min at 94°C followed by 35 cycles of 30 sec at 94°C for denaturation, 30 sec at hybridization temperatures based on the primers, and 30 sec at 72°C for extension; and then a final 7-min extension phase at 72°C.

The PCR products were purified using QIAquick PCR Purification Kit (Qiagen), according to the manufacturer’s instructions, using 50 μl of elution solution. Each eluent was evaporated and concentrated on a DNA SpeedVac (Thermo Scientific). The concentrated DNA solutions were assessed by NanoDrop 2000 (Thermo Scientific), and the DNA concentration was adjusted to 10 mg/mL using double-distilled water.

The oligonucleotide sequences used for amplification; and the length, G+C content, and number of CpG motifs contained in the amplicon are shown in Suppl. Table 1. The draft genome sequence of *L.lactis* LC-Plasma was available to the public (38) and was used for the design of primers and other purposes.

### Bone marrow (BM)-derived DC cultures

Four to 8-week-old female BALB/c wild-type mice were purchased from CLEA Japan. All animal care and experimental procedures were performed in accordance with the guidelines of the Committee for Animal Experimentation at Kirin Company. These studies were approved by the Committee for Animal Experiment at Kirin Company.

Flt3L-induced DCs were generated as follows. BM cells were extracted from BALB/c mice, and erythrocytes were removed by brief exposure to 0.168 M NH_4_Cl. Cells were cultured at a density of 5 × 10^5^ cells/mL for 7 days in RPMI 1640 medium (Life Technologies) containing 1 mM sodium pyruvate (Life Technologies), 2.5 mM HEPES (Life Technologies), 100 U/mL penicillin/100 μg/mL streptomycin (Life Technologies), 50 μM 2-ME (Life Technologies), 10% fetal calf serum (Life Technologies), and 100 ng/mL Flt3L (R&D Systems).

### Stimulating assay for pDCs

BM-derived DC cultures were stimulated with purified PCR products at a final concentration of 2 μg/mL in the presence of FuGENE^®^ HD Transfection Reagent (Promega) according to the manufacturer’s instructions. Briefly, FuGENE HD was added to the RPMI 1640 medium with 1000-fold dilution in final. Then, purified PCR products were added and the mixture was incubated for 5 min at room temperature. Each incubated mixture (50 μL) was added to 500 μL of culture medium containing BM-derived DCs at a density of 5.0 × 10^5^ cells/mL. For the experiment using oligomers (less than 50bp nucledotides), FuGENE^®^ HD was not used. After overnight incubation at 37°C in an atmosphere containing 5% CO_2_ and 95% air, the cell cultures were collected and centrifuged to obtain culture supernatants. The supernatants were stored at -80°C until analysis. IFN-α concentration was measured using the VeriKine^TM^ IFN-α ELISA Kit (PBL Assay Science), according to the manufacturer’s instructions.

### *In silico* analysis of bacterial genomes

*In silico* analysis was performed using Genetyx ver.9 software (GENETYX). We searched for 5’-purine-purine-CG-pyrimidine-pyrimidine-3’ (5’-RRCGYY-3’) and 5’-purine-TCG-pyrimidine-pyrimidine-3’ (5’-RTCGYY-3’), and the. total number of CpG hexamers (5’-NNCGNN-3’) in each genome was also calculated. In addition, we searched for 4 immunostimulatory motifs that were previously identified in LAB, as follows: BL07 (5’-GCGTCGGTTTCGGTGCTCAC-3’) (28), OL-LB7 (5’-CGGCACGCTCACGATTCTTG-3’) (29), ID35 (5’-ACTTTCGTTTTCTGCGTCAA-3’) (30), and OL-LG10 (5’-ATTTTTAC-3’) (31).

Genomic regions with low G+C content (e.g. G+C < 40%) were extracted using the source code that we created, based on the Perl Programming Language. When any 200 bp fragment was calculated with a G+C content ≥ 40%, genomic regions containing that fragment were designated as high-G+C regions. The genome sequence data of *Lactococcus lactis* susp. *lactis* LC-Plasma, *Leuconostoc mesenteroides* NBRC 100496 (synonym of ATCC 8293), *Lactobacillus acidophilus* NCFM, *Lactobacillus plantarum* WCFS1, *Lactobacillus casei* ATCC 334, *Lactobacillus fermentum* IFO 3956, *Lactobacillus rhamnosus* ATCC 53103, and *Pediococcus pentosaceus* ATCC 25745 were obtained from GENBANK and were used for *in silico* analysis.

